# Post-ictal neuronal network remodeling and Wnt pathway dysregulation in the intra-hippocampal kainate mouse model of temporal lobe epilepsy

**DOI:** 10.1101/604975

**Authors:** Kunal Gupta, Eric Schnell

**Affiliations:** Department of Neurosurgery, OHSU, Portland, Oregon, United States of America; Department of Anesthesiology and Perioperative Medicine, OHSU, Portland, Oregon, United States of America; VA Portland Health Care System, Portland, Oregon, United States of America

## Abstract

Mouse models of mesial temporal lobe epilepsy recapitulate aspects of human epilepsy, which is characterized by neuronal network remodeling in the hippocampal dentate gyrus. Observational studies suggest that this remodeling is associated with altered Wnt pathway signaling, although this has not been experimentally examined. We used the well-characterized mouse intrahippocampal kainate model of temporal lobe epilepsy to examine associations between post-seizure hippocampal neurogenesis and altered Wnt signaling. Tissue was analyzed using immunohistochemistry and confocal microscopy, and transcriptome analysis was performed on RNA extracted from anatomically micro-dissected dentate gyri. Seizures increased neurogenesis and dendritic arborization of newborn hippocampal dentate granule cells in peri-ictal regions, and decreased neurogenesis in the ictal zone, 2-weeks after kainate injection. Interestingly, administration of the novel canonical Wnt pathway inhibitor XAV939 daily for 2-weeks after kainate injection further increased dendritic arborization in peri-ictal regions after seizure, without an effect on baseline neurogenesis in control animals. Transcriptome analysis of dentate gyri demonstrated significant canonical Wnt gene dysregulation in kainate-injected mice across all regions for Wnt3, 5a and 9a. Intriguingly, certain Wnt genes demonstrated differential patterns of dysregulation between the ictal and peri-ictal zones, most notably Wnt5B, 7B and DKK-1. Together, these results demonstrate regional variation in Wnt pathway dysregulation in the early post-ictal period, and surprisingly, suggest that some Wnt-mediated effects might actually temper aberrant neurogenesis after seizures. The Wnt pathway may therefore provide suitable targets for novel therapies that prevent network remodeling and the development of epileptic foci in high-risk patients.

## Introduction

Mesial temporal lobe epilepsy can develop in humans after a number of neurologic insults, including trauma [1], infection [2], stroke [3] and febrile seizures [4]. Anti-epileptic drugs (AEDs) in these conditions can reduce seizure occurrence, however up to 40% of patients with epilepsy are considered medically refractory [5]. AEDs have been tested in a preventative fashion in post-traumatic epilepsy and effectively reduce early seizures after injury [6]; however, no treatment exists to prevent the development of delayed epilepsy after neurologic insults. Thus, an understanding of neuronal circuit remodeling after neurologic insults is necessary to optimally design preventative treatments.

Rodent models recapitulate key hallmarks of human temporal lobe epilepsy, including mesial temporal sclerosis, gliosis, mossy fiber sprouting and loss of hippocampal pyramidal and hilar neurons, and have contributed greatly to our understanding of mechanisms underlying epileptogenesis [7, 8]. In many of these models, an initial seizure is triggered using either focal (intraparenchymal) or global (systemic) manipulations, and is followed by a latent period during which rodents develop spontaneous recurrent seizures, similar to clinical observations in a subset of human patients [9].

It is increasingly recognized that circuit changes in epilepsy involve more than just the ictal zone where seizure activity initiates, and that peri-ictal regions are also remodeled to alter seizure threshold, creating secondary foci and wider epileptic networks [10, 11]. These peri-ictal regions and subsequent epileptic networks may provide additional therapeutic targets in the treatment of clinical epilepsy [12]. Consistent with a potentially wider epileptogenic zone, unilateral intrahippocampal kainate injection causes epilepsy characterized by bilateral hippocampal seizures in mouse models [11, 13], and is associated with increased neurogenesis in both the contralateral hippocampus and distant ipsilateral hippocampus [13, 14]. The contribution of these distant changes to epileptogenesis remains unknown, and the underlying transcriptional and molecular mechanisms that initiate these changes both in the ictal onset zone and the wider peri-ictal epileptogenic zone are poorly defined.

Recent work has begun to describe early transcriptional changes in both rodent models of epilepsy and human clinical samples, in hopes of identifying potential effectors of circuit remodeling during epileptogenesis. One potential effector system involves the Wnt (wingless/integrated) signaling pathway, which encompasses a large family of 19 secreted Wnt protein ligands, which bind a family of 10 membrane frizzled receptors and co-receptors. These regulate downstream canonical and non-canonical pathways, including the planar cell polarity and calcium pathways [15]. These signals have been implicated in neurogenesis and dendrite formation in the adult rodent hippocampus [16, 17], and dysfunction of these processes are associated with epileptogenesis [18–20]. Dysregulation of hippocampal Wnt and mTor (mechanistic target of rapamycin) pathways was observed using microarray analysis in a hypoxic neonatal mouse seizure model; in this study, beta-catenin was dysregulated in an activity-dependent manner independent of the hypoxic insult, suggesting that canonical Wnt signaling may play an early role in epileptogenesis [21].

The association between Wnts and epileptogenesis is only beginning to be explored and may offer novel therapeutic targets that prevent the development of epilepsy after neurologic insults [22]. Here, we investigate hippocampal Wnt signaling changes during epileptogenesis, using the mouse intra-hippocampal kainate model of epilepsy. First, we characterize seizure-induced alterations in hippocampal neurogenesis after kainate using the *POMC-EGFP* transgenic mouse, in which newly born dentate granule cells transiently express eGFP (enhanced green fluorescent protein) [23]. We subsequently evaluate the role of the Wnt pathway in these early changes by administration of XAV939, a canonical Wnt antagonist [24], and transcriptionally profile the whole dentate gyrus to characterize changes in Wnt gene transcription during early epileptogenesis.

## Materials and methods

### Animal husbandry

Experiments were performed utilizing the *-13/+8POMC-EGFP* transgenic mouse line (MGI: 3776091), in which newborn dentate granule cells are labeled with eGFP for up to 2 weeks after birth [23]. Homozygous and heterozygous male mice were studied between 3-4 months of age, as female mice have been shown to have neither a latent period nor hippocampal discharges with intrahippocampal kainate [25]. Mice were housed according to local IACUC guidelines, with food and water *ad libitum*. All procedures and animal handling were performed in accordance with the *Guide for the care and use of laboratory animals* and were approved by the Oregon Health & Science University Institutional Animal Care and Use Committee (IACUC).

### Stereotactic intrahippocampal kainate injection

After induction with isoflurane by spontaneous respiration, mice were anesthetized with isoflurane by nose cone, head-shaved and secured in the stereotactic apparatus with ear bars. The scalp was sterilized with betadine and topical lidocaine gel was applied for local anesthesia. A single midline sagittal incision was performed with a #10 scalpel blade and the bregma was visualized under stereoscopic magnification. Drill coordinates were acquired relative to bregma. A single burr hole was placed at X +1.8, Y −2.1 mm and debris was cleared with sterile saline irrigation. The injection needle was slowly inserted to target, Z −1.7 mm from the dura. Sterile normal saline vehicle or kainate (Cayman Chemicals, 0.5mg/ml in normal saline) were delivered by Hamilton syringe connected to a Quintessential Stereotaxic Injector (Stoelting); 100µl were injected over 1 minute. The needle was left in place for 2 minutes to prevent reflux of the injection, and then slowly withdrawn. The skin was closed with dermal glue (Vetbond, 3M) and the mouse allowed to recover in a warmed chamber. Seizures were scored by a modified Racine scale for 2 hours after injection; stages 1 and 2 demonstrated freezing, mastication and head nodding, stage 3 demonstrated forelimb clonus, stage 4 demonstrated rearing, stage 5 demonstrated rearing and falling, stage 6 demonstrated “popcorn” type seizures [26, 27]. Only mice that were observed to undergo at least one seizure measuring 3-6 on the Racine scale were included for study in the kainate group [28]. After recovery, mice were provided food saturated with diluted pediatric acetaminophen (3.2mg/ml) for 24 hours and assisted feeding with soft food daily until sacrifice.

### Drug preparation and intra-peritoneal injection

XAV939 (Millipore Sigma) was dissolved in filter-sterilized 10% DMSO / 45% saline / 45% poly-ethylene glycol-400 (Affymetrix) at a final concentration of 1mg/ml. Animals received intraperitoneal injections daily for 14 days with vehicle or XAV939 (5 mg/kg) after recovery from surgery.

### Immunohistochemistry

Mouse brains were retrieved after terminal anesthesia and trans-cardiac perfusion with 4% paraformaldehyde, post-fixed overnight, embedded in 2% agarose and sectioned at 100µm thickness on a vibratome. Slices were blocked and permeabilized in PBS containing 5% goat serum and 0.4% Triton-X for 1 hour. Primary antibodies included anti-GFP (A-21311 polyclonal rabbit antibody, Alexafluor 488 pre-conjugated, Thermo Fisher Scientific), anti-c-Fos (2250, monoclonal rabbit antibody, Cell Signaling Technology) and anti-ZnT3 (197 004, polyclonal guinea pig antibody, Synaptic Systems Germany), which were applied at 1:500 dilution overnight at 4°C; Alexafluor-conjugated secondary antibodies (donkey isotype, Thermo Fisher Scientific) were applied at 1:1000 dilution overnight at 4°C. DAPI was applied at 1:10,000 dilution for 15 minutes and sections were mounted on slides with Fluoromount-G (Thermo Fisher Scientific).

### Immunohistochemical analysis

Immunohistochemically stained sections were imaged with confocal microscopy (Carl Zeiss LSM 780, Jena, Germany). Four to five animals were used in each experimental group. For dendritic arbor and migration analysis, images were acquired using a 20x objective. Two sections per region per mouse were analyzed; image stacks were 22.0µm thick with 1.1µm steps. eGFP+ cell counts were performed manually in ImageJ using image stacks of 23.4 µm thickness using a 10x objective, from two sections for each location in each animal. Dorsal and ventral regions were identified anatomically. For contralateral dentate gyrus, all parameters measured were similar between both contralateral dorsal and ventral hippocampus, therefore data and figures are displayed for dorsal contralateral hippocampal sections. An eGFP+ cell was considered within the granule cell layer if the center of its soma was between 10µm below the inferior border of the granule cell layer (to include the subgranular zone) to the outer granule cell layer border. Granule cell layer width and volume were measured using ImageJ (NIH). Dendritic arbors of eGFP+ newborn dentate granule cells were measured using FilamentTracer (Imaris BitPlane, Zurich, Switzerland). Mean arbor length per cell was calculated by dividing the total arbor length for a given section by the number of eGFP+ cells within that section. Migration analysis was performed manually in ImageJ (NIH), by measuring the distance of the eGFP+ cell body from the hilus-subgranular zone border. Data were subjected to statistical analysis and graphs prepared using Prism (GraphPad, San Diego, CA USA). Data are presented as mean ± standard error.

### Transcriptional analysis

Bilateral dentate gyri were anatomically microdissected in an RNAse-free environment. The ipsilateral dentate was hemisected into dorsal and ventral parts, the contralateral dentate was processed whole. Tissue was then placed in Qiazol (Qiagen), macerated with an RNAse-free pestle (Kimble Chase) and stored at −80°C until processing. RNA was isolated using the Universal Plus mini Kit (Qiagen) with QIAcube automation. RNA quality assessment was performed using the Agilent 2100 Bioanalyzer with a Eukaryote total RNA Nano chip. All samples received an RNA integrity score of >9. Reverse transcription (RT) was performed using the SuperScript VILO cDNA synthesis kit (Life Technologies) with 650ng of input RNA per 80μl reaction. Following reverse transcription, 2µl of cDNA was used in the PCR reaction with 10µl TaqMan universal master mix and 1µl of 20x gene specific TaqMan assay in a total volume of 20µl and loaded onto the QuantStudio instrument. The qPCR assays were performed on the QuantStudio Real-time PCR System (Life Technologies) using a single master-mix per TaqMan probe set, for Wnt3, Wnt5A, Wnt5B, Wnt7A, Wnt7B, Wnt8A, Wnt8B, Wnt9A, WLS (wnt ligand secretor) and DKK-1 using TUBA1A as the endogenous control. Four 384-well plates were used; each plate was setup to contain a complete biologic group in order to minimize variation. Additionally, a calibrator sample (pooled RNA) was used in each plate assayed, to ensure concordance between plates. Data were collected using Applied Biosystems QuantStudio™ 12K Flex Software v1.2.2. Fold change in expression were calculated by ΔΔCt method and analyzed using a two-tailed Students’ t-test; regional comparisons were performed by 2-way ANOVA with repeated measures.

## Results

### Unilateral intra-hippocampal kainate injection causes early bilateral hippocampal activation and delayed regional differences in dentate histology

We used the intrahippocampal kainate injection model of temporal lobe epilepsy to study post-ictal neurogenesis in the hippocampus of adult male *POMC-EGFP* mice. As previously described [29], unilateral injection of kainate into the hippocampal CA1 region elicited modified Racine stage 3-6 seizures in mice [26], including behavioral freezing, prolonged mastication, clonic movements of the forelimbs and hindlimbs, rearing, and “popcorn” type seizures. In comparison, unilateral hippocampal injection of saline (vehicle control) had no effects on behavior or seizures. After kainate or saline injection, mice were allowed to recover and maintained for up to 2 weeks prior to analysis (Fig 1A).

**Fig 1.**
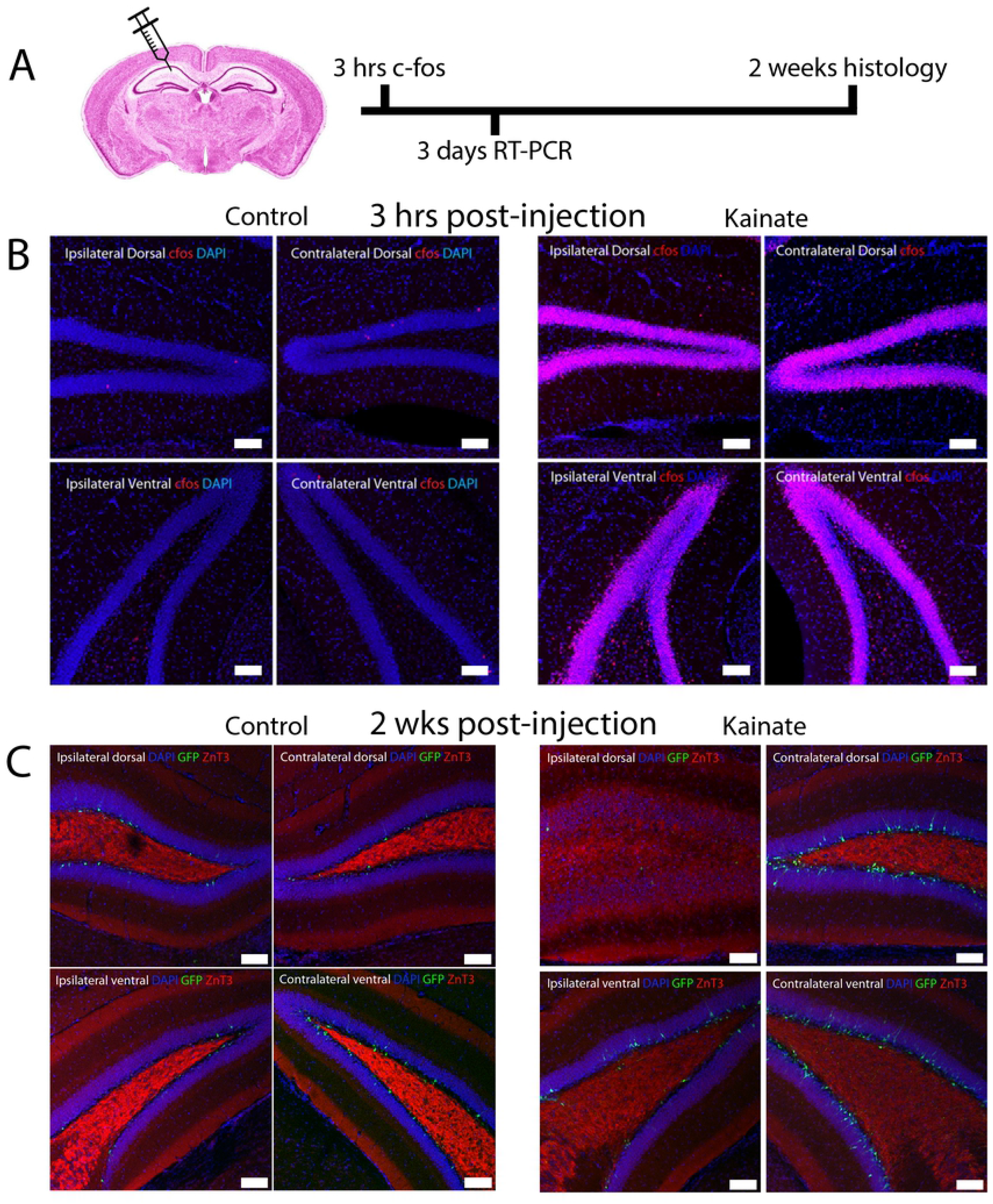
Intrahippocampal kainate injection causes widespread bilateral dentate granule cell activation, followed by delayed focal mossy fiber sprouting. (A) Experimental timeline after intrahippocampal injection. (B) c-Fos expression (red) in the dentate gyri from hippocampal quadrants 3hrs after saline (control; left) or kainate (seizure; right) injection into the ipsilateral/dorsal region CA1. Images show extensive bilateral c-Fos expression 3hrs after kainate in the dentate gyrus. (C) The mossy fiber terminal marker ZnT3 is localized to the denate hilus as expected 2wks after saline injection (control; left), and localizes to the granule cell and molecular layers adjacent to the site of kainate injection in post-ictal *POMC-EGFP* animals. Scale bar 100µm.

We first examined expression of c-Fos, a marker of neuronal activation, 3 hours after kainate injection. When compared with saline-injected controls, unilateral kainate injection caused diffuse and widespread c-Fos expression throughout both the ipsilateral and contralateral dentate gyrus, in both the dorsal and ventral hippocampus (Fig 1B). Thus, although kainate was injected unilaterally, there was diffuse bilateral activation of neurons in the hippocampal dentate gyrus, indicating generalized seizure activity that propagated to both hemispheres.

The hippocampus was examined 2 weeks after intrahippocampal injection to determine the delayed effects of focal kainate injection on the histology of the dentate gyrus (Fig 1C). We defined the ictal zone as dentate gyrus near the site of kainate injection (ipsilateral dorsal hippocampus), and compared it with neighboring peri-ictal regions in both the ipsilateral (ipsilateral ventral) and contralateral dentate gyrus. At this 2-week timepoint, we observed marked differences in histology between the dentate gyrus adjacent to the site of injection, the ictal zone, and the neighboring peri-ictal zones.

We observed granule cell dispersion, a hallmark of temporal lobe epilepsy [29–31], only in the ictal zone but not in peri-ictal regions (Fig 1C). Similarly, mossy fiber sprouting into the granule cell and molecular layers of the dentate gyrus, another finding strongly associated with epilepsy, was also only observed in the ictal region (Fig 1C). Thus, despite robust bilateral neuronal activation induced by kainate activation, subsequent histologic changes induced by seizure activity differed between the ictal and peri-ictal zones. This suggested differences in the molecular and cellular signaling events between these regions, which could relate to the differential development of histological changes post-seizure induction as well as peri-ictal epileptogenic foci.

### The novel Wnt antagonist XAV939 does not quantitatively alter neurogenesis in control animals

Wnt signaling controls neuronal migration during development as well as dendritic outgrowth from immature neurons [15]. Global Wnt pathway dysregulation has been demonstrated after seizures [21, 32], and there is growing interest in the mechanisms by which specific Wnt pathway mediators may contribute to epileptogenesis [22]. As post-ictal neuronal remodeling has been implicated in the development of epilepsy [33–35], we hypothesized that changes in Wnt signaling after seizures might alter neuronal circuit remodeling in the dentate gyrus. We administered XAV939, a novel small molecule canonical Wnt antagonist [24], daily by intra-peritoneal injection for 2 weeks to determine whether this affects neuronal remodeling after seizures. XAV939 inhibits tankyrase 1 and 2, which then leads to beta-catenin degradation and blockade of downstream canonical Wnt pathway signaling [24].

First, we examined the effects of canonical Wnt inhibition on the early development of adult-born dentate granule cells in saline injected (control) *POMC-EGFP* animals treated with vehicle or XAV939 for 2 weeks (Fig 2A-D), to determine whether canonical Wnt signaling plays a role in the early phase of constitutive neurogenesis. *POMC-EGFP* mice express eGFP in immature adult-born hippocampal granule cells, and can be used to both quantitatively and qualitatively assess adult neurogenesis [23]. Based on our analysis of eGFP-expressing immature adult born granule cells, XAV939 treatment in control animals did not significantly change adult-born dentate granule cell dendrite arbor length (Fig 2B), cell count (Fig 2C) or cell migration (Fig 2D). This suggests that these constitutive processes are not dependent upon intact canonical Wnt signaling pathways under baseline conditions.

**Fig 2.**
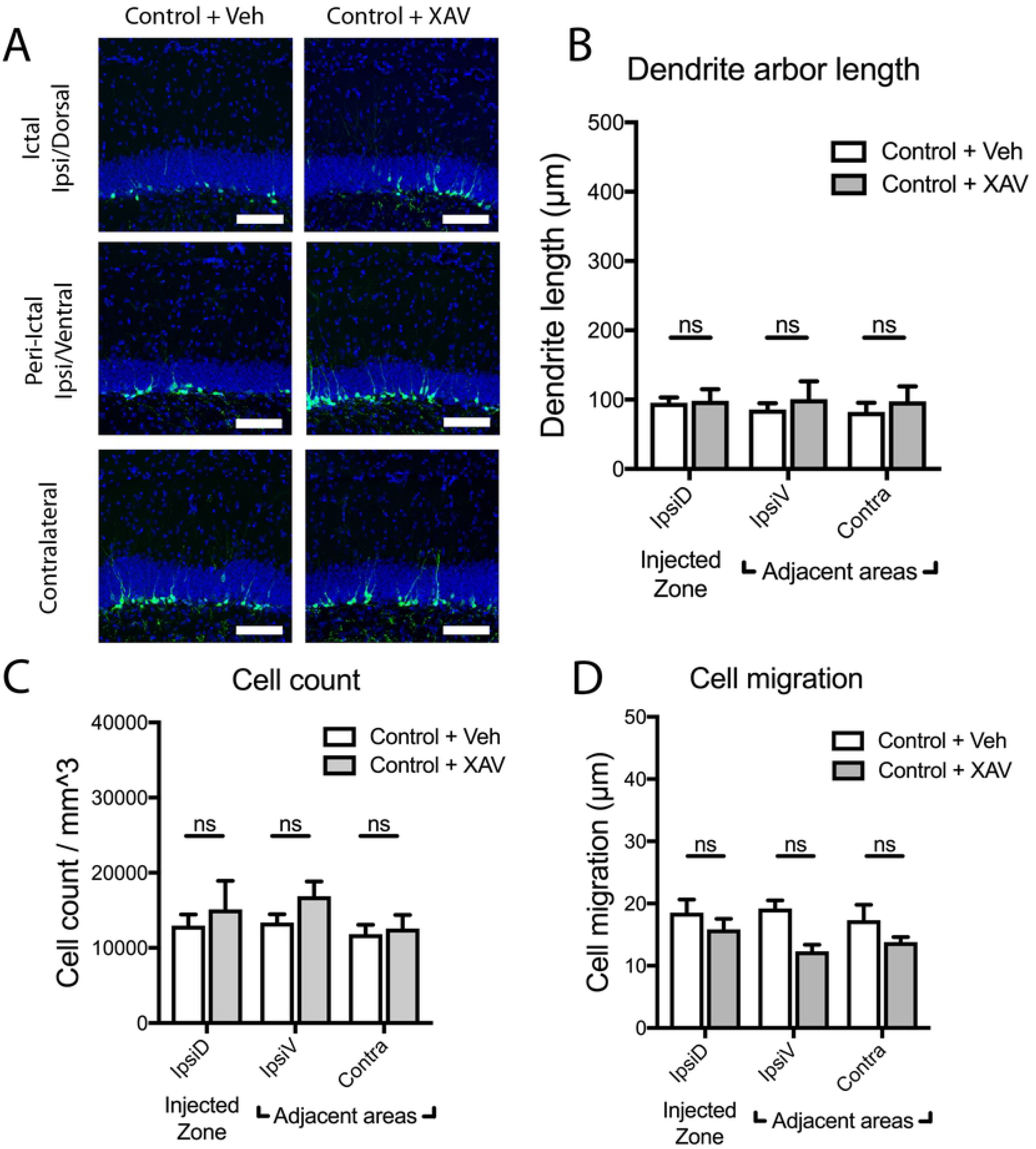
Treatment with novel Wnt antagonist XAV939 does not alter newborn dentate granule cell maturation in control animals. (A) Images demonstrate eGFP+ newborn dentate granule cells two weeks after intrahippocampal saline injection (control) in the ipsilateral dorsal, ipsilateral ventral, and contralateral dentate gyrus, of mice treated with vehicle vs. XAV939 daily for two weeks after injection. Scale bars 100µm. (B-D) In saline-injected control animals, continuous Wnt inhibition by XAV939 treatment does not alter newborn dentate granule cell arbor length (B), cell density (C), or cell migration (D), when compared with vehicle-treated control mice (ns = not significant).

We subsequently examined whether Wnt inhibition by XAV939 affected gross structural changes in the dentate gyrus after kainate-induced seizures. As previously observed [29–31], seizure induction by kainate markedly increased dentate granule cell layer dispersion in the ictal zone (Fig 3A). Ictal zone granule cell layer dispersion was unaffected by XAV939 treatment (Fig 3B), indicating that it occurs independent of canonical Wnt signaling. Granule cell dispersion was not observed in peri-ictal regions after seizures; and in these regions, granule cell layer thickness was also not affected by XAV939 treatment (Fig 3B). Importantly, XAV939 did not drive granule cell layer dispersion in any region in control/saline treated animals (Fig 3B), suggesting that constitutive canonical Wnt signaling is not needed to maintain granule cell layer organization in healthy mice.

**Fig 3.**
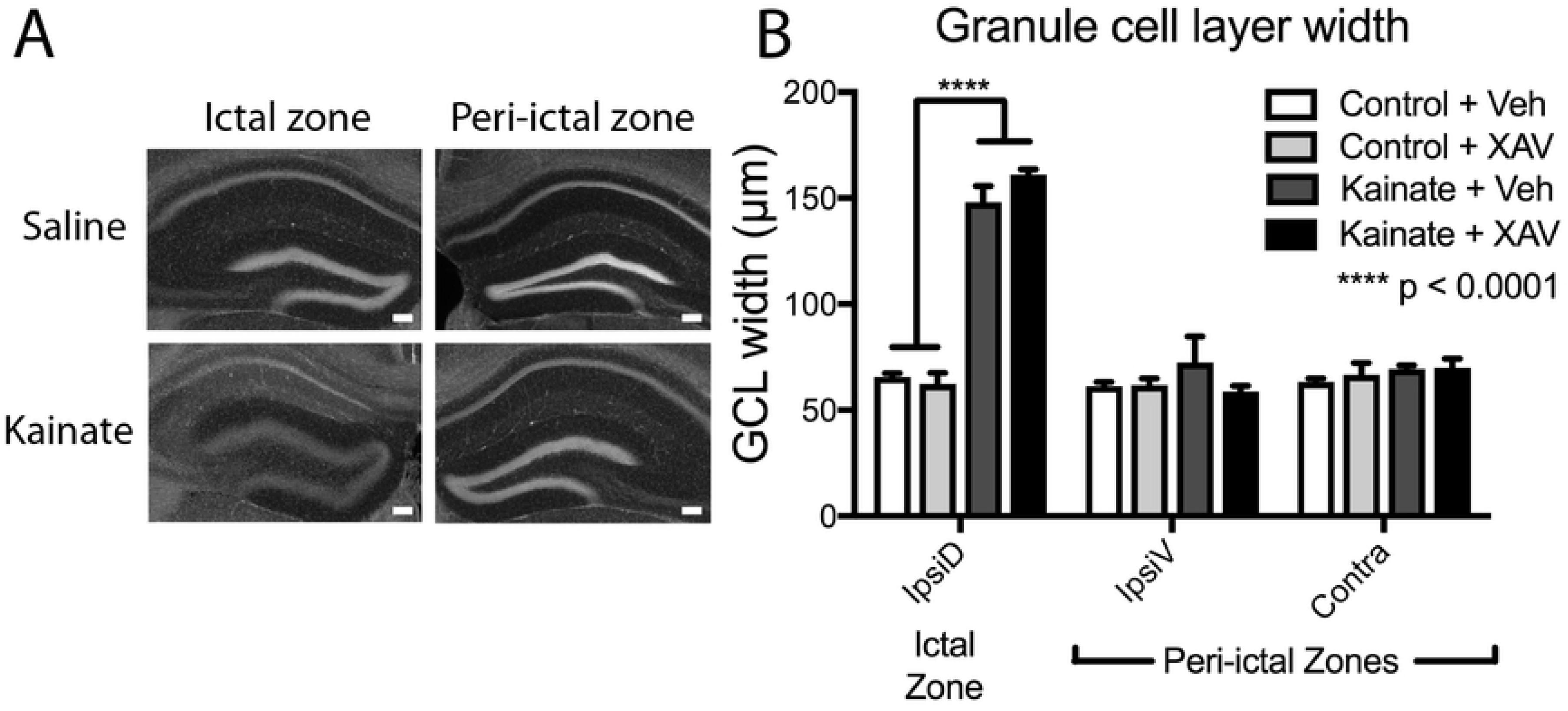
Granule cell dispersion in the ictal zone is not affected by XAV939 treatment. (A). DAPI-stained images of coronal whole dentate gyrus after saline (control) and kainate injection in the ictal (injected) and peri-ictal (non-injected) regions of the dentate gyrus. Scale bar 100µm. (B). Dentate granule cell layer dispersion is only seen in the ictal zone after seizure induction by kainate, and this was not affected by daily administration of Wnt antagonist XAV939 for two weeks after kainate.

### XAV939 increases dendrite growth by newborn neurons in peri-ictal regions after seizure

Seizures drive hippocampal neuronal remodeling in a variety of mouse models [33, 35], and these changes may relate to the subsequent development of epilepsy [28, 34, 35]. We therefore investigated whether seizure induction was associated with region-specific changes in the morphologic maturation of dentate granule cells born after seizure. We used *POMC-EGFP* mice to allow for both quantitative and morphologic assessment of newborn dentate granule cells, as they express eGFP for up to 2-weeks post-mitosis [23].

Prior work using the pilocarpine model of epilepsy in mice demonstrated that seizures dramatically increased dendrite arbor length of newborn granule cells in the molecular layer of the dentate gyrus [28]. However, in the intrahippocampal kainate model, we observed increased dendritic arbor length of newborn dentate granule cells only in the peri-ictal regions, and not in the ictal region (Fig 4A, B). Interestingly, the seizure-induced dendrite growth was very similar in both the ventral ipsilateral hippocampus and the contralateral hippocampus. As this peri-ictal dendrite growth closely resembled the phenotype observed in the pilocarpine model, we believe that close proximity to the ictal (kainate injection) zone produced a region-specific effect on the dentate in this model that was qualitatively different than that which occurred in the remainder of the hippocampus.

**Fig 4.**
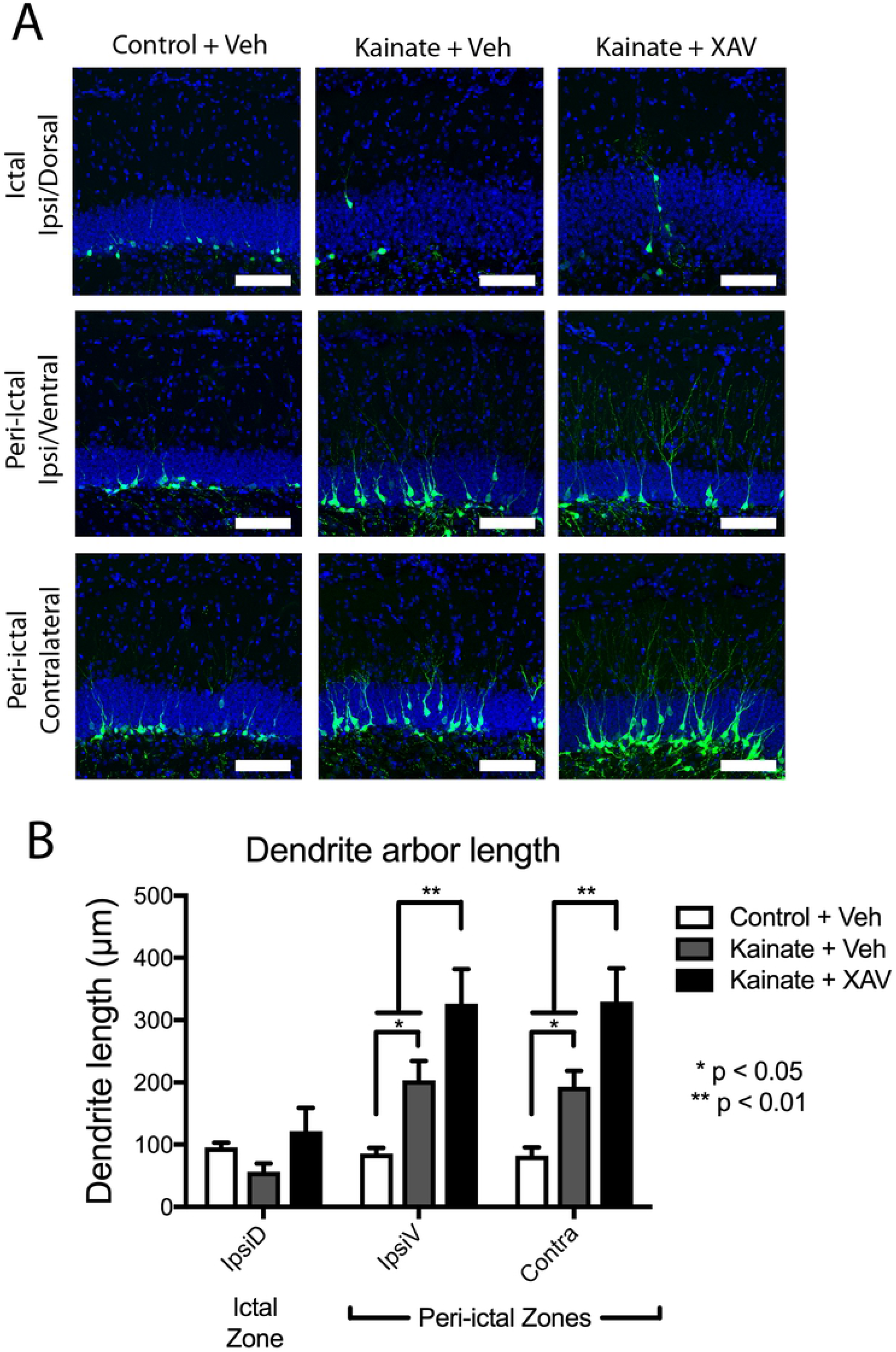
Post-ictal increase in newborn granule cell dendrite length is augmented by Wnt inhibition. (A) Images of *POMC-EGFP*+ adult-born dentate granule cell neurons from indicated hippocampal regions two weeks after intrahippocampal kainate or saline injection, followed by daily systemic treatment with XAV939 vs. vehicle control. Scale bars 100µm. (B) In peri-ictal regions, mean dendritic arbor length per eGFP+ cell increased significantly after kainate-induced seizure in peri-ictal zones, and was further increased by XAV939 treatment.

Wnt antagonism had an unanticipated effect on the growth of dentate granule cell dendrites after seizures, as XAV939 treatment significantly increased growth of dendrites by newly born cells in the peri-ictal regions after seizures (Fig 4A, B). This was not a general effect of XAV939 on dendritic arborization, as it had no effect on dendritic branching in control/saline treated animals (Fig 2A, B). This suggests that the effects of canonical Wnt signaling on neuronal remodeling primarily manifested after seizures, and that Wnt signaling in the peri-ictal dentate gyrus actually restricts aberrant growth of dendrites from granule cells after seizures.

### XAV939 modulates post-ictal dentate neurogenesis and neuronal migration

Consistent with previous reports [13, 36, 37], intrahippocampal kainate injection significantly decreased the density of newborn dentate granule cells in the ipsilateral dorsal dentate gyrus / ictal zone, adjacent to the site of kainate injection (Fig 5A, B). This decreased neurogenesis in the ictal zone was not altered by XAV939 treatment (Fib 5A, B). However, kainate-induced seizures did not increase neurogenesis in the contralateral or ipsilateral ventral regions (Fig 5A, B), unlike previously reported [13]. XAV939 treatment increased the cell count in the ipsilateral peri-ictal region after kainate-induced seizure, which was not observed in the contralateral dentate gyrus (Fig 5B).

**Fig 5.**
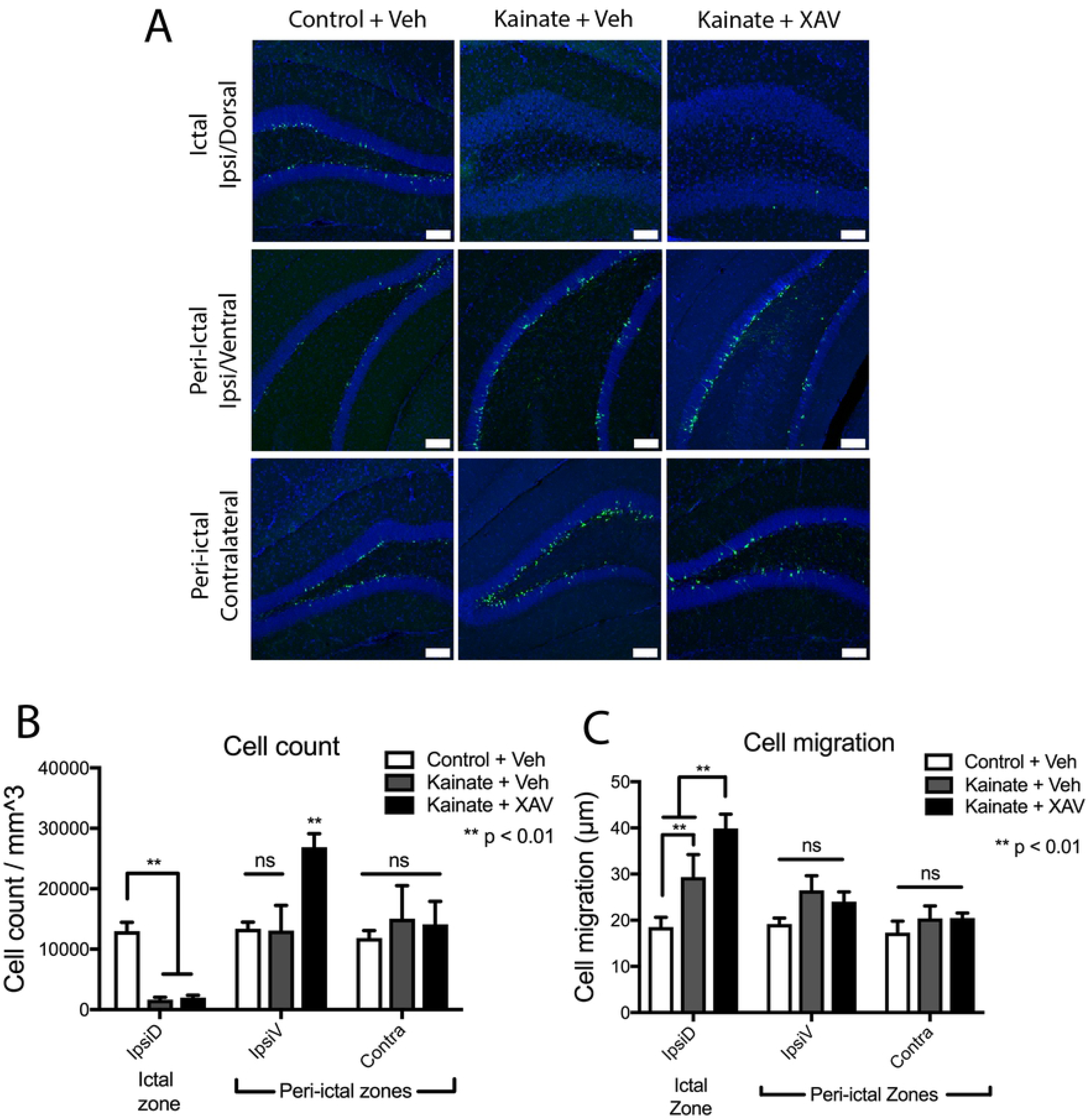
Modulation of neurogenesis and cell migration by the Wnt antagonism. (A) *POMC-EGFP*+ newborn dentate granule cells 2wks after kainate-induced seizure, followed by vehicle or XAV939 treatment, in ictal and peri-ictal regions. Scale bars 100µm. (B) Newly born (*POMC-EGFP*+) granule cell density decreased in the ictal zone, and was not rescued by XAV939 treatment. In the ipsilateral peri-ictal region, cell density increased after XAV939 treatment only in kainate-treated mice. No XAV939-induced change in cell density was observed in the contralateral dentate. (C) Newborn dentate granule cell migration increased in the ictal zone 2wks after seizure induction, which was further increased by XAV939 treatment. Cell migration was unchanged after kainate in the peri-ictal zones.

Kainate-induced seizures significantly increased the migration of immature neurons from the subgranular zone in the ictal, but not peri-ictal, regions (Fig 5A, C). Again, XAV939 treatment after kainate had region-specific effects, such that XAV939 increased neuronal migration in the ictal region, whereas neuronal migration was unaffected by XAV939 in the peri-ictal regions (Fig 5A, C). This suggests that canonical Wnt signaling after kainate-induced seizures again appeared to primarily normalize aberrant phenotypes after seizures, as inhibition of this pathway led to a more dramatic phenotype.

### Transcriptional profiling of ictal and peri-ictal dentate gyri

To characterize the changes in Wnt signals that might be involved in post-ictal dentate gyrus remodeling, we performed a transcriptomic analysis of candidate Wnt molecules from the dentate gyri of epileptic mice, based on prior reports of their involvement in neurogenesis, neuronal morphogenesis or dentate specific expression in the Allen Brain Atlas (Allen Institute for Brain Science) [32, 38-40]. Dentate gyri were anatomically micro-dissected from mice that experienced intrahippocampal kainate-induced seizures, and from control mice that had received intrahippocampal saline injection (Fig 6A-B). To help determine how Wnt signals might differentially be associated with the profoundly different structural phenotypes between ictal and peri-ictal regions, we subdivided dentate gyri anatomically, corresponding to the ipsilateral dorsal dentate gyrus at the injection site (ictal zone), and 2 peri-ictal regions, the ipsilateral ventral dentate and contralateral dentate. Profiling was performed via quantitative RT-PCR of tissue 3-days after kainate vs. saline injection, to determine whether kainate-induced seizure altered expression of any of these genes at a timepoint that would be expected to affect the development of dentate granule cells born after seizure induction. Transcriptional data for each individual gene and region are reported in supplementary table S1, comparative data for each gene are demonstrated in Fig 6C.

**Fig 6.**
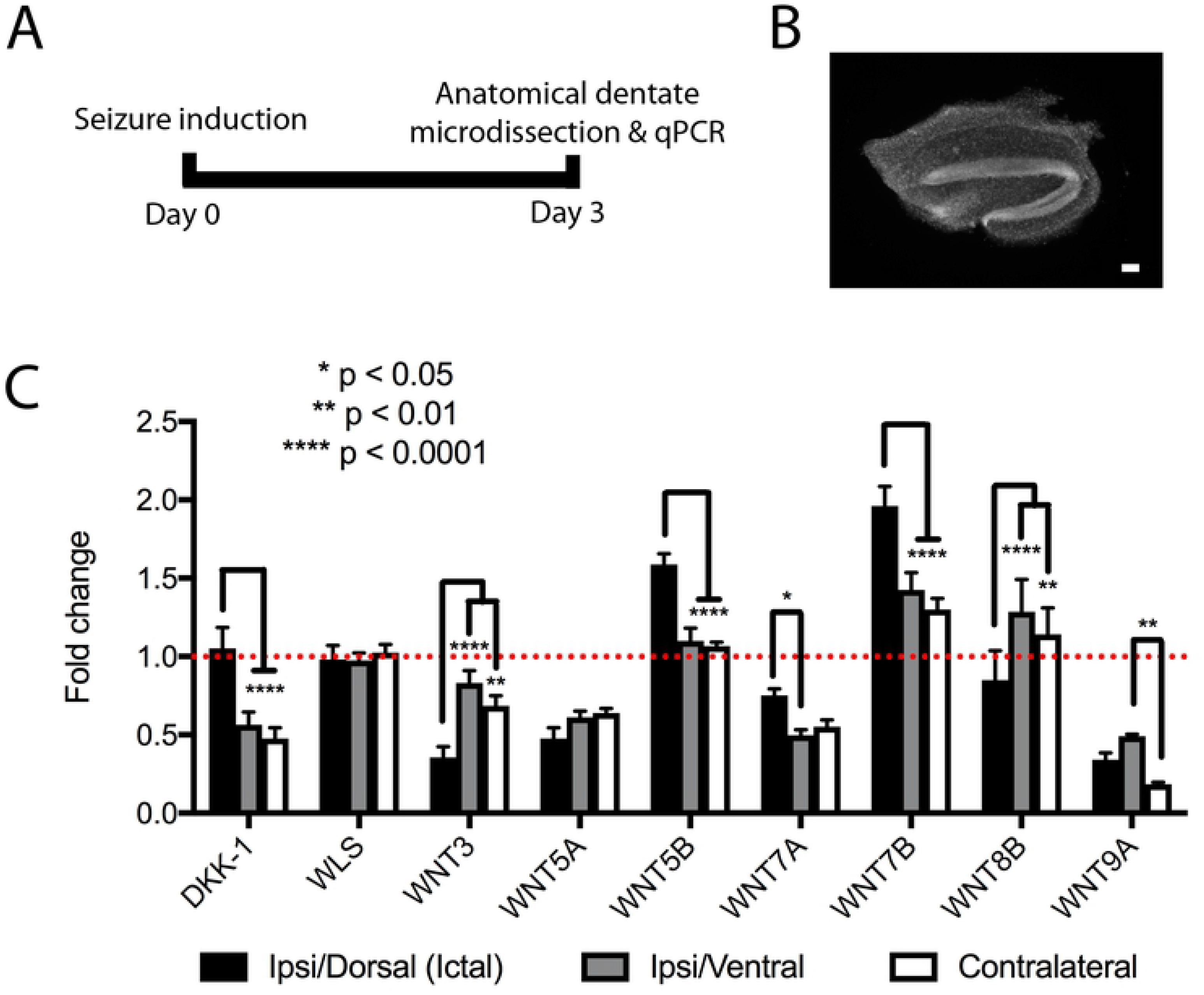
Transcriptional profiling of the whole dentate gyrus 3 days after seizure induction. (A) Timeline of transcriptional analysis. (B) Image of a representative DAPI-labeled cross-section of micro-dissected dentate gyrus used for transcriptional analysis. Scale bar 100µm. (C) Relative Wnt pathway gene transcription in the ictal and peri-ictal regions 3d after seizure induction. Various patterns are observed, whereby certain genes are selectively upregulated in the ictal zone (Wnt5b and Wnt7b), selectively downregulated in ictal (Wnt3) or peri-ictal regions (Dkk-1), or relatively unchanged from baseline (WLS, Wnt8b).

In the ictal zone, the largest changes in transcriptional regulation were seen in Wnt3 and Wnt7b, whereas in the peri-ictal zones, the largest changes in transcriptional regulation were seen in DKK-1 and Wnt7a (Fig 6C). Interestingly, changes in individual Wnt gene expression in both peri-ictal regions showed striking similarities in their patterns of dysregulation, such that these genes were either upregulated or downregulated in both peri-ictal regions, and often in direct contrast to the transcriptional changes (or lack thereof) in the ictal region (e.g., DKK-1, Wnt5b, Fig 6C). This pattern matches the distinct phenotypes noted between ictal and peri-ictal regions (Figs 3-5), as well as the striking similarity between the ipsilateral/ventral and contralateral dentate gyri. Certain genes were similar in all regions after seizure, being either unchanged from control (WLS and Wnt8b), or downregulated (Wnt5a and Wnt9a, supplementary table 1), but the majority of Wnt genes analyzed showed similar differential expression patterns between ictal and peri-ictal (ipsilateral/ventral and contralateral) regions (Fig 6C).

## Discussion

It is increasingly recognized that not only are remote regions of the dentate gyrus and hippocampus affected by focal seizures, but that seizure-related alterations in the neuronal circuit connectivity of these regions might also contribute to brain hyperexcitability [11, 41]. For instance, in epileptic humans, epileptic foci are associated with structural changes and reduced seizure thresholds in neighboring brain, and these neighboring regions potentially act as independent epileptogenic regions [42–44]. Using the *POMC-EGFP* mouse to visualize granule cells born after an acute seizure event, we demonstrate that structural abnormalities occur in peri-ictal hippocampus after seizures, which include changes in newborn neuron migration and dendrite arbor formation, and that these post-ictal structural changes are affected by Wnt modulation.

### Modulation of post-seizure hippocampal remodeling by Wnt signaling

Aberrant circuit rewiring in the hippocampal dentate gyrus is a hallmark of temporal lobe epilepsy in humans, and may contribute to the formation of epileptic foci [10, 31]. This rewiring includes altered granule cell migration, axonal arborization, and dendritic morphology, much of which involves qualitative and quantitative changes in post-seizure neurogenesis [18, 28, 33-35]. Two weeks after focal kainate-induced seizures, we observed marked heterogeneity in cellular responses between different regions of the hippocampus. These included marked granule cell dispersion and decreased neurogenesis in the dentate gyrus adjacent to the site of kainate injection (ictal zone), and upregulation of dendritic growth in the peri-ictal regions distant from the injection site. These findings are consistent with prior observations, which demonstrated similar differential effects on neurogenesis between ictal and peri-ictal zones using BrdU-based mitotic cell labeling and cell filling [13].

In addition to increased overall granule cell dispersion in the ictal zone [29, 45], migration of newly born dentate eGFP+ granule cells also increased in the ictal zone 2 weeks after kainate-induced seizures (Fig 5). As Wnts control the migration of neuroblasts during development [17, 46, 47], we had hypothesized altered Wnt signaling may contribute to this accelerated migration. Surprisingly, however, we observed that migration of newly born dentate granule cells in the ictal zone increased with pharmacologic antagonism of canonical Wnt signaling using the novel Wnt antagonist XAV939 [24]. Thus, our data are more consistent with a model in which canonical Wnt signaling actually inhibits the aberrant migration of newly generated granule cells, and that some other signal (or loss thereof, for instance, reelin [48]) causes aberrant newborn granule cell migration during epileptogenesis.

Interestingly, canonical Wnt antagonism by XAV939 also increased the length of newborn dentate granule cell dendrites after seizure, specifically in the peri-ictal regions. Again, in this region, Wnt signaling after seizures appeared to counteract some signal which was driving the increased dendritic outgrowth. We base this interpretation on the observation that XAV939 did not accelerate dendritic outgrowth in control conditions (intrahippocampal control saline injections), indicating that canonical Wnt signaling is not required for dendritic outgrowth during constitutive neurogenesis. Given prior evidence that specific Wnt signals control dendritic morphogenesis during neuronal development [39, 49, 50], we were surprised that canonical Wnt antagonism did not noticeably affect constitutively-generated adult-born granule cells under control conditions. This suggests that Wnts may play differential roles for granule cells generated during early neonatal development from those generated in adults.

Under control conditions, Wnt antagonism with XAV939 did not affect neurogenesis. Constitutive beta-catenin activation has previously been shown to mediate expansion of Tbr2+ intermediate neurons [46], and lentivirus-mediated Wnt3 transduction has been shown to mediate expansion of DCX+ neurons in the dentate gyrus in 15wk old rats [17]. Our findings that small chemical Wnt antagonism did not affect baseline neurogenesis appear inconsistent with this, however other compensatory mechanisms may be present.

In contrast to prior BrdU-labeling studies after intrahippocampal kainate, we also did not observe an increased number of eGFP+ cells in the peri-ictal regions [35]. It is not clear what caused this discrepancy, but perhaps since eGFP+ cells represent the aggregate population of newly born granule cells generated during the preceding 2 weeks [23], the lack of change may reflect the overall integrated sum of post-ictal neurogenesis over time, encompassing both the initial increase and subsequent decrease in neurogenesis rates that both occur within this two week window [35].

Although we did not observe effects of canonical Wnt inhibition on constitutive neurogenesis, our analysis was limited to morphological features of adult-born granule cells during their early maturation, and does not preclude Wnt modulation of later stages of neuronal maturation. Similarly, our data also do not rule out Wnt-mediated control of neurogenesis via alternate downstream Wnt signaling pathways, such as the calcium or planar cell polarity pathways, or Wnt-mediated effects on circuit function that do not involve neurogenesis. Finally, although XAV939 has been shown to be selective for the Wnt pathway and to not affect the CRE (cAMP response element), NF-kB (nuclear factor kappa-light-chain-enhancer of activated B cells), or TGFb (transforming growth factor beta) pathways in reporter lines [24], and was effective in both *in vivo* and *in vitro* assays involving mice and neural cell types at the doses similar to this study [51–53], unexpected off-target drug effects must still be considered.

### Transcriptional changes in Wnt mediators after seizure

We examined transcriptional changes in the dentate gyrus in the early post-ictal period and found dysregulated expression of key Wnt genes known to modulate neuronal growth and architecture [17, 38, 39, 54-56]. Perhaps the most striking finding in the analysis of these transcriptional changes involves how the parallel histologic changes in peri-ictal and contralateral remodeling were accompanied by similar transcriptional patterns within Wnt pathway genes, both of which were very distinct from transcription in the ictal zone.

In terms of specific genes, Wnt3 was downregulated in the ictal zone and unchanged in peri-ictal regions. As Wnt3 positively regulates hippocampal neurogenesis, as well as spinal cord neurogenesis and neurite outgrowth [17, 56], its downregulation in the ictal zone is consistent with our observation of reduced neurogenesis in this region, and suggests that post-ictal downregulation of Wnt3 might contribute to the reduction of neurogenesis in the zone. Wnt7b is expressed primarily in the dentate gyrus of the hippocampus during the course of early post-natal development, and appears to drive dendritic growth and branching of hippocampal neurons via Dvl (disheveled segment polarity protein homolog) and Rac1 (Ras-relate C3 botulinum toxin substrate 1) [39]. We observed an overall upregulation of Wnt7b in the ictal and peri-ictal zones, which is consistent with a possible contribution of Wnt7b to enhanced post-ictal dendrite growth. In this case, however, as the canonical Wnt antagonist XAV939 did not prevent, but actually enhanced, dendritic outgrowth, any potential roles of Wnt7b in driving dendritic outgrowth after seizures would likely be mediated by a non-canonical downstream Wnt pathway.

Our transcriptional characterization of Wnt signaling changes during early epileptogenesis, however, does not clearly implicate a specific gene, and it does not exclude multiple Wnt pathways acting in concert. Additionally, we performed our profiling at 3 days after kainate to analyze changes in Wnt genes during early epileptogenesis, but Wnt pathways will need to be examined further in a detailed fashion using specific Wnt gene modulation in specific hippocampal cell types at serial time-points to obtain a more complete picture of dynamic changes in Wnt signals that might occur over longer timeframes. Additionally, although the pattern of associations between structural phenotypes and Wnt signaling suggest that these changes may be related, we are unable to make causative association between individual Wnt genes and any specific outcomes. Future studies involving manipulations of specific Wnt signals and mediators will hopefully allow us to ascribe specific functional roles of individual candidate molecules to specific post-ictal changes in dentate structure, and eventually, function.

## Conclusions

The intrahippocampal kainate model is a well-established model of temporal lobe epilepsy, in which kainate-injected mice manifest spontaneous seizures after several weeks [9]. Wnt antagonism with XAV939 altered the course of post-ictal dentate remodeling, however it remains to be seen how this drug, or Wnt signal modulation in general, would affect the eventual development and severity of subsequent spontaneous seizures. Future studies with XAV939, and eventually with more targeted approaches specific to individual Wnts, will allow us to extend our data to determine how Wnts might be involved in epileptogenesis. Better understanding of Wnt pathway dysregulation in epilepsy may identify therapeutic targets that in high-risk human patients prevent the development of focal epilepsy in response to inciting conditions such as trauma, tumor, infection and others.

## Acknowledgements

This material was supported in part by the Department of Veterans Affairs Merit Review Award I01-BX002949 (ES), a Department of Defense CDMRP Award W81XWH-18-1-0598 (ES), a Neurosurgery Research and Education Foundation Fellowship (KG) and P30NS061800 (OHSU) awards. We thank Dr. Gary Westbrook of the Vollum Institute for comments on our manuscript, Dr. Stefanie Kaech-Petrie of the OHSU Advanced Light Microscopy Core for assistance with imaging, and research assistant Sarah Mader for technical support. The contents of this manuscript do not represent the views of the U.S. Department of Veterans Affairs or the United States Government.

